# ALDH2 DEFICIENCY INCREASES SUSCEPTIBILITY TO BINGE ALCOHOL-INDUCED GUT LEAKINESS, ENDOTOXEMIA, AND ACUTE LIVER INJURY IN MICE THROUGH THE GUT-LIVER AXIS

**DOI:** 10.1101/2022.04.23.489282

**Authors:** Wiramon Rungratanawanich, Xin Wang, Toshihiro Kawamoto, Saravana Babu Chidambaram, Byoung-Joon Song

## Abstract

Mitochondrial aldehyde dehydrogenase 2 (ALDH2) is the major enzyme responsible for metabolizing toxic acetaldehyde to acetate and acts as a protective or defensive protein against various disease states associated with alcohol used disorder (AUD), including alcoholic liver disease (ALD), and elevated oxidative stress. We hypothesized that *Aldh2*-knockout (KO) mice are susceptible to binge alcohol-mediated liver injury than wild-type (WT) mice through increased gut leakiness and endotoxemia. Therefore, this study aimed to investigate the protective role of ALDH2 in binge alcohol-induced gut permeability, endotoxemia, and acute inflammatory liver injury by exposing *Aldh2*-KO or WT mice to a single oral dose of binge alcohol 3.5, 4.0, or 5.0 g/kg. Our findings showed for the first time that ALDH2 deficiency in *Aldh2*-KO mice increases their sensitivity to alcohol-induced oxidative and nitrative stress, enterocyte apoptosis, and nitration of gut tight junction (TJ) and adherent junction (AJ) proteins, leading to their degradation. These resulted in gut leakiness and endotoxemia in *Aldh2*-KO mice after exposure to a single dose of ethanol even at 3.5 g/kg, while no changes were observed in the corresponding WT mice. The elevated serum endotoxin (lipopolysaccharide, LPS) and/or bacterial translocation contributed to systemic inflammation, hepatocyte apoptosis, and subsequently acute liver injury, indicating the disruption in the gut-liver axis. Furthermore, treatment with Daidzin, an ALDH2 inhibitor, exacerbated ethanol-induced cell permeability and reduced TJ/AJ proteins in T84 human colonic cells. These changes were reversed by Alda-1, an ALDH2 activator, indicating a crucial role of ALDH2 in protecting against alcohol-induced epithelial barrier dysfunction. All these findings suggest that *ALDH2* deficiency or gene mutation in humans is a risk factor to alcohol-mediated gut and liver injury, and ALDH2 could be an important therapeutic target against alcohol-associated tissue/organ damage.

**Highlights:**

- Binge alcohol increases oxidative and nitrative stress in the intestine and liver.
- Binge alcohol causes gut leakiness, endotoxemia, and acute liver injury.
- Leaky gut is caused by elevated degradation of nitrated intestinal TJ/AJ proteins.
- *Aldh2*-KO mice are susceptible to binge-alcohol-induced leaky gut and liver injury.
- ALDH2 inhibition increases alcohol-induced T84 colonic epithelial cell permeability.

## 1. Introduction

Excessive alcohol (ethanol) drinking is known to cause more than 200 diseases and injuries worldwide [1]. Binge alcohol consumption, which represents greater than 75% of alcohol-associated social and economic loss, leads to neurobehavioral abnormality and functional injury to multiple major organs such as the intestine, liver, and brain [2]. In the gastrointestinal (GI) tract, heavy alcohol drinking is frequently associated with small intestinal bacterial overgrowth (SIBO) and gut leakiness with translocation of bacteria and bacterial-derived products such as endotoxin (e.g., lipopolysaccharide or LPS) into the circulation. Endotoxin specifically LPS is a significant pathogen-associated molecular pattern (PAMP) molecule produced from gram-negative bacteria. LPS can directly stimulate immune cells in the gut lamina propria, contributing to local inflammation. It can also increase oxidative and nitrative stress (or nitroxidative stress), which stimulates post-translational modifications (PTMs) and degradation of intestinal tight junction (TJ) and adherent junction (AJ) proteins, resulting in gut leakiness [3–10]. The elevated gut leakiness further allows translocation of endotoxin (LPS) and bacteria into the liver through the enterohepatic circulation, causing inflammatory and fatty liver [11, 12]. In fact, gut leakiness is a common feature observed in many people with alcohol used disorder (AUD), including alcoholic liver disease (ALD) with inflammation [3, 11]. The increased serum endotoxin (endotoxemia) is also shown to positively correlate with the levels of alcohol consumption and advanced liver diseases such as fibrosis and cirrhosis [13]. Additionally, alcohol-associated gut leakiness with endotoxemia has been observed in experimental models of mice [9, 14] and rats [5, 9] as well as people with alcohol intoxication [8], demonstrating a common phenomenon conserved among different species.

In oxidative alcohol metabolism, mitochondrial aldehyde dehydrogenase 2 (ALDH2) is the major enzyme responsible for metabolizing toxic acetaldehyde to acetate, which is eventually degraded to CO2 and H2O [15]. ALDH2 is also involved in detoxification of highly reactive lipid aldehydes, such as malondialdehyde (MDA) and 4-hydroxynonenal (4-HNE) under oxidative stress conditions and acts as a protective or defensive protein against multiple disease states associated with AUD (e.g., ALD) [6, 16–18]. More than 560 million people, particularly East Asians, carry a dominant negative *ALDH2* gene mutation (*ALDH2*2*) and have very low ALDH2 activity. These people show typical, physiological responses (e.g., facial flushing, palpitations, nausea, and tachycardia) and increased susceptibility to alcohol-associated tissue injury and vulnerability to cancer (i.e., oral and gastrointestinal cancer) due to alcohol drinking [19–23]. In mice, deletion of *Aldh2* gene or *Aldh2* knockout (KO) contributes to increased sensitivity to alcohol-related DNA damage and tissue injury compared with wild-type (WT) mice [24–28]. However, the underlying mechanism by which ALDH2 deficiency causes susceptibility to alcohol-mediated organ damage, especially after binge alcohol exposure, remains to be elucidated. In particular, the contributing role of gut leakiness in promoting acute liver injury via the gut-liver axis and the detailed mechanisms have not been systematically studied in *Aldh2*-KO mice. Here, we hypothesized that *Aldh2*-KO mice are more susceptible to binge alcohol-mediated inflammatory liver injury than the corresponding WT mice through increased gut permeability and endotoxemia. Therefore, this study aimed to investigate the protective role of ALDH2 in binge alcohol-induced gut leakiness, endotoxemia, and acute liver injury in *Aldh2*-KO and WT mice after exposure to a single oral dose of alcohol 3.5, 4.0, or 5.0 g/kg. In addition, the cultured T84 human colonic cells were used to strengthen our mouse results and conduct mechanistic studies on the protective role of ALDH2 in binge alcohol-induced intestinal barrier dysfunction and hepatotoxicity.

## 2. Materials and methods

All animal experimental procedures were carried out by following the National Institutes of Health (NIH) guidelines for small animal experiments and approved by the NIAAA Institutional Animal Care and Use Committee. All mice were maintained under controlled environment: temperature (22±3°C), humidity (45-55%) and artificial lighting (12-h light/dark cycle) with free access to food and water provided *ad libitum*. Young male, age-matched inbred global *Aldh2*-KO mice on C57BL/6J background [24, 25] and WT mice (n ≥ 3~4/group) were fasted overnight before they were orally exposed to a single dose of binge alcohol at 3.5, 4.0, or 5.0 g/kg. Control mice were orally administrated a vehicle (water). The intestinal enterocytes, blood, and liver tissue were collected at 1 or 6 hours after the ethanol exposure for observation of intestinal or liver injury, respective, as previously reported [9, 29, 30], and stored at −80 ^o^C until further characterization.

### 2.1 Histological analysis

Sections of fresh liver and intestine from alcohol-exposed or control mice were fixed in 10% neutral-buffered formalin for histopathological analysis. Paraffin-embedded blocks of formalin-fixed individual liver and intestine (ileum near the cecum) sections were cut at 4 μm and stained with hematoxylin and eosin (H&E) by American Histolabs, Inc. (Gaithersburg, MD, USA).

### 2.2 Terminal deoxynucleotidyl transferase dUTP nick end labeling (TUNEL) assay

Formalin-fixed liver and intestinal samples were processed, and 4-μm thick paraffin sections were used. The ApopTag Peroxidase *in situ* apoptosis detection kit (Millipore, Billerica, MA, USA) was employed to assess DNA strand breaks to evaluate the rates of apoptotic cell death of hepatocytes and enterocytes, respectively, by the TUNEL analysis.

### 2.3 Serum alanine aminotransferase (ALT) and aspartate aminotransferase (AST)

Serum ALT and AST levels in each animal were determined by using the colorimetric assay kit (TECO Diagnostics, Anaheim, CA, USA) according to the manufacturer’s instruction.

### 2.4 Serum endotoxin and reactive oxygen species (ROS)

Serum endotoxin (LPS) concentrations were measured by the quantitative endpoint chromogenic Limulus amebocyte lysate (LAL) assay (Lonza Inc., Walkersville, MD, USA) with a concentration range of 0.1-1 endotoxin units per milliliter (EU/mL). Serum ROS levels were determined by using Amplex® Red reagent 10-acetyl-3,7-dihydroxyphenoxazine (Thermo Fisher Scientific) in combination of horseradish peroxidase (HRP) to detect hydrogen peroxide (H2O2) or peroxidase activity and produce the red-fluorescent oxidation product, resorufin, with excitation and emission of approximately 530 nm and 590 nm, respectively, as described [31–34].

### 2.5 Measurements of hepatic lipid peroxidation, interleukin-6 (IL-6), and TNFα

Hepatic lipid peroxides MDA and 4-hydroxyalkenal (μM) were assessed by the reaction of N-methyl-2-phenylindole at 45°C to yield a stable chromophore product with maximal absorbance at 586 nm using the Lipid Peroxidation Microplate Assay Kit (Oxford Biomedical Research, Oxford, MI, USA), as recently described [35]. The levels of hepatic IL-6 and TNFα were quantified by using the IL-6 (mouse) and TNFα (mouse) ELISA kits (BioVision, Milpitas, CA, USA) following the manufacturer’s instructions.

### 2.6 Measurements of hepatic and intestinal caspase-3 activity

Hepatic and intestinal caspase-3 activity was measured by the Caspase-3 assay kit (Abcam, Cambridge, UK) with the liver and gut lysates, respectively. The absorption at 400 nm was recorded with a microplate reader and used to calculate the caspase-3 activity by comparing with the standard contained in the kit.

### 2.7 Immunoblot analysis

The enterocytes of the small intestine from each animal were homogenized with RIPA buffer to prepare gut extracts. Protein concentrations were determined via the Bradford protein assay, and equal amounts of protein from different samples were separated by 10% SDS/PAGE and transferred to nitrocellulose membranes. These membranes were probed with the respective primary antibody against ZO-1 (1:1,000 dilution; Abcam), Occludin (1:1,000 dilution; Abcam), Claudin-1 (1:1,000 dilution; Abcam), Claudin-3 (1:1,000 dilution; Abcam), Claudin-4 (1:1,000 dilution; Abcam), β-catenin (1:1,000 dilution; Abcam), E-cadherin (1:1,000 dilution; Cell Signaling), Plakoglobin (1:200 dilution; Santa Cruz), ALDH2 (1:1,000 dilution; Abcam), CYP2E1 (1:1,000 dilution; Abcam), iNOS (1:1,000 dilution; Abcam), Bax (1:1,000 dilution; Cell Signaling), Cleaved (active)-caspase-3 (1:1,000 dilution; Cell Signaling), or GAPDH (1:5,000 dilution; Cell Signaling). Horseradish peroxidase (HRP)-conjugated goat anti-rabbit or anti-mouse IgG (1:5,000 dilution; Cell Signaling) was used as the secondary antibody. Relative protein images were assessed by enhanced chemiluminescence (ECL) substrates, and their immunoreactive band intensities were quantified by densitometry using ImageJ software (National Institutes of Health).

### 2.8 Immunoprecipitation

Immunoprecipitation was performed with the gut homogenates, and the total 0.5 mg of proteins were equally pooled from individual WT or *Aldh2*-KO mice of control or alcohol-exposed groups. The lysates were incubated with the specific antibody against each indicated target protein, including β-catenin, Occludin, or Claudin-1 (Abcam, Cambridge, UK). Pierce™ protein A/G magnetic beads (Thermo Fisher Scientific™, USA) were used by following the manufacturer’s protocol. The immunoprecipitated proteins were subsequently analyzed by immunoblot analysis using the specific antibody against 3-nitrotyrosine (3-NT) and determined by ECL substrates, as previously described [9, 10].

### 2.9 16S metagenomic sequencing and bioinformatics

Stool samples were collected from the cecum of each mouse and rapidly frozen at −80 °C. DNA was extracted using Mag-Bind Universal Pathogen DNA Kit (Omega Bio-Tek, Norcoss, GA) following the manufacturer’s protocols. DNA sequencings for the bacterial 16S ribosomal V3-V4 region of each stool sample were performed at the Omega Bioservices Inc. (www.omegabioservices.com). Sequence data was processed with Qiime 2 against the Greengenes 13.8 database to assign taxonomy.

### 2.10 Cell culture and FITC-Dextran 4-kDa analysis for measurement of intestinal cell permeability

Human T84 colonic carcinoma cells were grown on collagen-coated polycarbonate membrane Transwell® inserts with a surface area of 0.33 cm^2^ and 0.4 μm pore size (Sigma-Aldrich, St. Louis, MO) in a humidified incubator at 37 °C under 5% CO2 in DMEM/F-12 medium supplemented with 5% fetal bovine serum (FBS), 1% non-essential amino acids solution, and 1% antibiotic-antimycotic solution, as described [9, 10]. After 7-14 days, confluent monolayer cells were observed, and these cells were treated with ethanol (130-150 mM, for ~1.5 h) [36] in the absence or presence of pretreatment with an ALDH2 inhibitor (Daidzin, 10 μM, for 12 h) [37, 38] or an ALDH2 activator (Alda-1, 20 μM, for 12 h) [39–41]. Fluorescein isothiocyanate-labeled 4-kDa dextran (FITC-D4; 1 mg/mL, Sigma-Aldrich) was added to the apical side of cells and incubated for 1 h at 37 °C. Then, the fluorescence intensity of FITC-D4 in the culture medium of basolateral side was measured with a microplate reader at excitation and emission spectra of 485 nm and 540 nm, respectively, to evaluate epithelial cell permeability. Data were reported as relative fluorescent units (% baseline).

### 2.11 ALDH2 activity and immunoblot analysis

T84 cells were grown in a humidified incubator at 37 °C under 5% CO2 for 7-14 days. After confluent monolayer formation, the cells were treated with 130-150 mM ethanol for ~1.5 h [36] in the presence or absence of an ALDH2 inhibitor (Daidzin) [37, 38] or an ALDH2 activator (Alda-1) [39–41] and harvested for further analyses. Protein concentrations were determined via the Bradford protein assay. Equal amounts of protein were separated by SDS/PAGE and transferred to nitrocellulose membranes followed by probing with the respective primary antibody against ALDH2 (1:1,000 dilution; Abcam), Occludin (1:1,000 dilution; Abcam), ZO-1 (1:1,000 dilution; Abcam), β-catenin (1:1,000 dilution; Abcam), or GAPDH (1:5,000 dilution; Cell Signaling). ALDH2 activity was evaluated by Mitochondrial Aldehyde Dehydrogenase (ALDH2) Activity Assay Kit (Abcam, Cambridge, UK) following the manufacturer’s protocol.

### 2.12 MTT assay analysis for measurement of intestinal cell viability

After 7-14 days of culture, T84 cells were pretreated with an ALDH2 inhibitor (Daidzin) or an ALDH2 activator (Alda-1) for 12 h followed by ethanol exposure 130-150 mM for ~1.5 h. After brief exposure to ethanol, cell medium was then replaced with MTT reagent (Abcam, Cambridge, UK), and the cells were incubated at 37°C for additional 3 h. MTT solution (Abcam, Cambridge, UK) was added, and the cells were further incubated at room temperature for 15 min before absorbance was measured with a microplate reader at OD 590 nm.

### 2.13 Immunofluorescence and confocal microscopy

T84 cells were plated onto chamber slides. The cells were then fixed with 4% paraformaldehyde for 15 min at room temperature and washed 3 times with PBS for 10 min each, followed by blocking with 3% (w/v) BSA solution in PBS for an additional 1 h. After removal of the blocking solution, the remaining cells were incubated with the specific antibody against ALDH2, Occludin, ZO-1, or β-catenin (Abcam, Cambridge, UK) at 4 °C overnight. Alexa Fluor 488-labeled goat anti-rabbit IgG (Thermo Fisher Scientific™, USA) was used as the secondary antibody for immunofluorescence detection. The T84 cells were then mounted with Fluoromount-G™ Mounting Medium (Thermo Fisher Scientific™, USA), and the fluorescence images were visualized by using a confocal microscope (Carl Zeiss, Oberkochen, Germany).

### 2.14 Statistical analysis

Statistical significance was determined by two-way or one-way ANOVA and Tukey’s multiple comparison post-test analysis to compare the means of multiple groups using Graph Pad Prism version 8.0 (Graph Pad Software, San Diego, CA, USA). Data are presented as means ± SD and considered statistically significant at p < 0.05.

## 3. Results

### 3.1 Binge alcohol exposure increased gut inflammation and apoptosis with elevated serum endotoxin and oxidative stress in Aldh2-KO mice

Histological H&E staining showed that a single low dose of ethanol (3.5 g/kg, p.o.) was sufficient to cause gut inflammation in *Aldh2*-KO mice after 1 h, while no inflammatory cells are present in the gut of the corresponding WT mice. At the higher doses of ethanol (4.0 or 5.0 g/kg, p.o.), *Aldh2*-KO mice exhibited deformation of the intestinal villi structure with blunt, short, and less villi density, compared with the corresponding WT mice. In contrast, inflammatory cells were present in the WT mice only after exposure to ethanol at 5.0 g/kg without clear structural alteration in the intestine (Fig. 1A). TUNEL assay for DNA strand breaks and apoptotic cell death revealed that a single low dose of ethanol (3.5 g/kg, p.o.) was sufficient to induce enterocyte apoptosis in *Aldh2*-KO mice after 1 h but not in the corresponding WT mice. Moreover, at the higher doses of ethanol (4.0 or 5.0 g/kg, p.o.), *Aldh2*-KO mice exhibited markedly elevated levels of apoptotic enterocytes compared with WT counterparts (Fig. 1B). These findings were also supported by a significant increase in caspase-3 activity, an apoptosis indicator, in the gut lysates of *Aldh2*-KO mice 1 h after alcohol exposure, while little changes were observed in the corresponding WT mice. In addition, significantly higher levels of serum endotoxin (LPS) and ROS were observed in alcohol-exposed *Aldh2*-KO mice, 1 h after alcohol exposure, compared to those in the control or the corresponding WT mice (Fig. 1C), suggesting a protective role of ALDH2 in alcohol-mediated intestinal injury.

**Figure 1:**
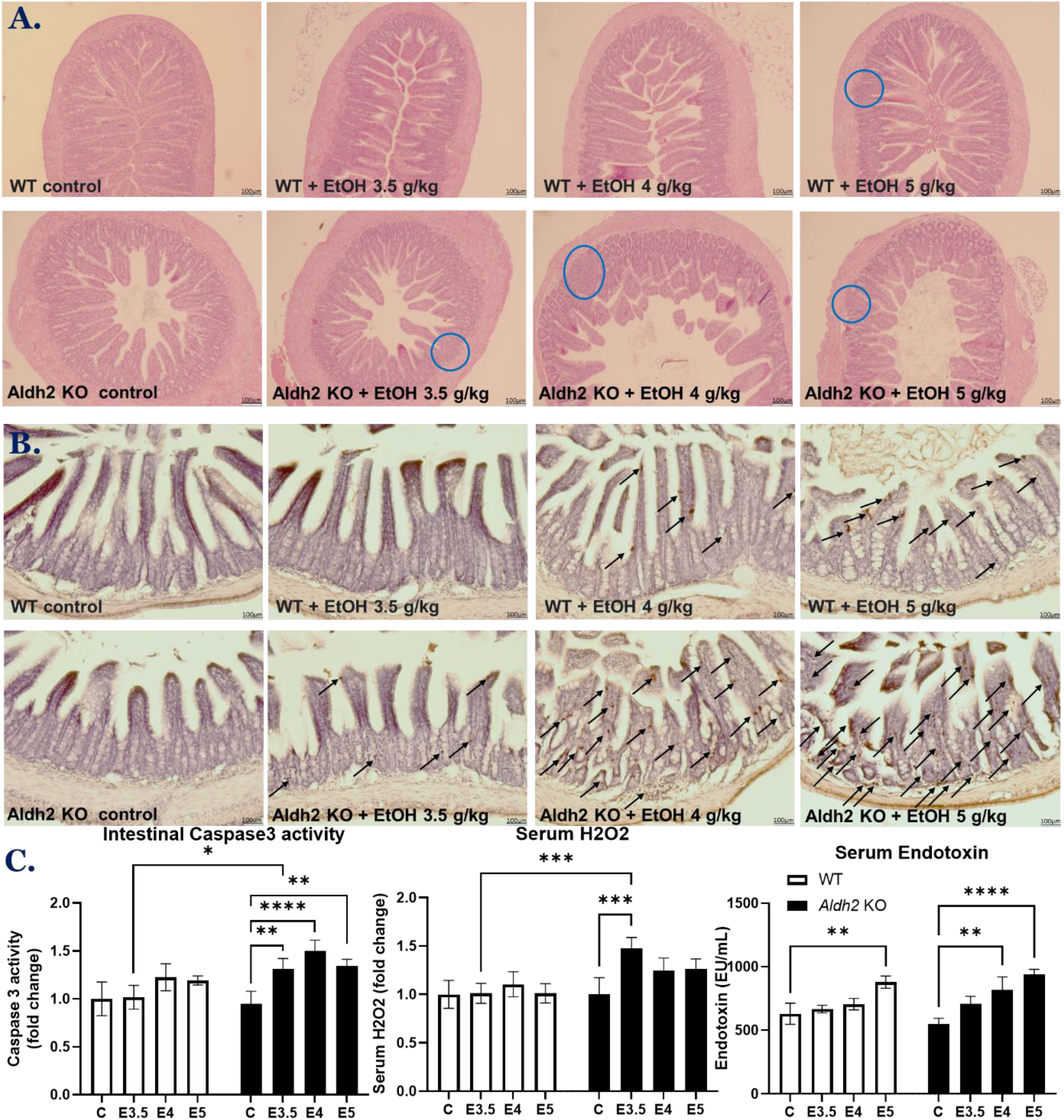
Binge alcohol exposure increased gut inflammation and apoptosis with elevated serum endotoxin and ROS in *Aldh2*-KO mice. (**A**) Representative H&E staining and (**B**) TUNEL staining of formalin-fixed small intestinal sections of WT and *Aldh2*-KO mice exposed to a single dose of ethanol 3.5, 4.0, or 5.0 g/kg (n ≥ 3~4/group), as indicated. The blue circles in (**A**) represent the area of inflammation and arrows in (**B**) indicate apoptotic cells. (**C**) The levels of intestinal caspase-3 activity, serum endotoxin, and serum ROS are presented for the indicated groups. CON: Control, E3.5: EtOH 3.5 g/kg, E4: EtOH 4.0 g/kg, E5: EtOH 5.0 g/kg. Bar scale, 100 μm in A and B. Data are expressed as means ± SD where statistical significance was determined using two-way ANOVA; * p<0.05, ** p<0.01, *** p<0.001, **** p<0.0001.

### 3.2 Binge alcohol exposure accelerated liver inflammation, oxidative stress, and apoptosis in Aldh2-KO mice

After verifying the impact of *Aldh2* deletion in elevated gut injury, we further investigated the effects of alcohol on the liver of *Aldh2*-KO compared with WT mice through the gut-liver axis. Similar to the intestine, histological H&E staining revealed that a single low dose of ethanol (3.5 g/kg, p.o.) caused inflammatory cell infiltration into the liver of *Aldh2*-KO mice after 6 h but not in the corresponding WT mice. Greater numbers of inflammatory cells were observed in the livers of *Aldh2*-KO mice after exposure to higher doses of ethanol (4.0 or 5.0 g/kg, p.o.), whereas only ethanol 5.0 g/kg, p.o. was able to enhance inflammatory cell infiltration into the liver of WT mice (Fig. 2A). These histological results were also supported by significant increases in the levels of pro-inflammatory cytokines IL-6, TNF-α, and lipid peroxides in the liver lysates of binge alcohol-exposed *Aldh2*-KO mice 1 h after alcohol exposure compared with those in the control and WT mice exposed to the same doses of ethanol (Fig. 2B). In addition, TUNEL staining showed that a single dose of ethanol 3.5 g/kg, p.o. caused hepatocyte apoptosis, similar to the enterocytes, in the *Aldh2*-KO mice 6 h after alcohol exposure but not in corresponding WT mice. Furthermore, the higher doses of ethanol (4.0 or 5.0 g/kg, p.o.) markedly increased the rates of hepatocyte apoptosis in *Aldh2*-KO mice compared with WT counterparts (Fig. 2C). Consistently, caspase-3 activity was significantly elevated in the liver lysates of binge alcohol-exposed *Aldh2*-KO mice 6 h after alcohol exposure compared with the corresponding WT mice. Serum AST and ALT levels were significantly elevated in *Aldh2*-KO mice exposed to a single dose of ethanol after 6 h, while no significant changes were observed in the corresponding WT mice (Fig. 2D), suggesting a protective role of ALDH2 in alcohol-mediated hepatic injury.

**Figure 2:**
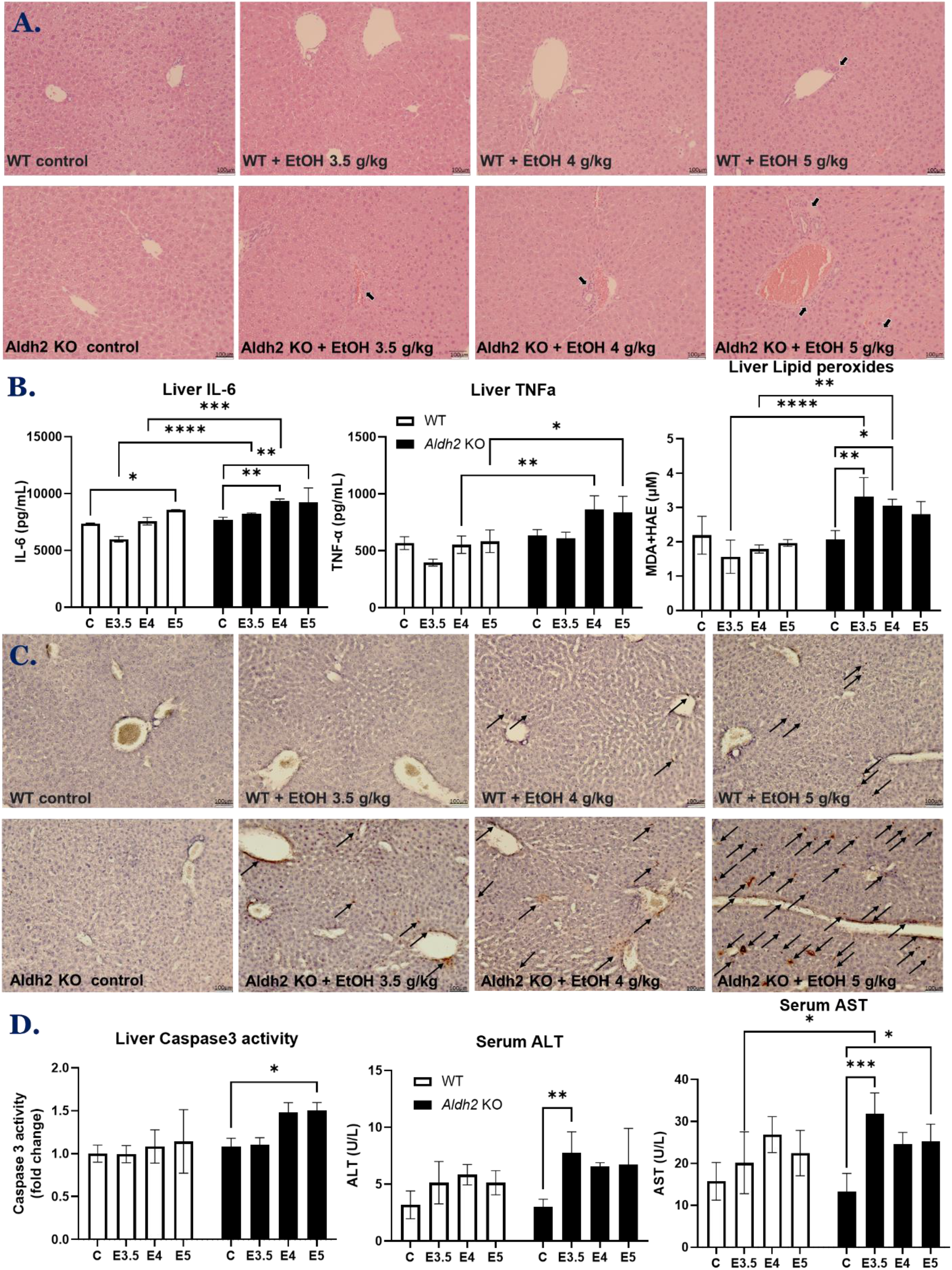
Binge alcohol exposure accelerated acute liver inflammation, apoptosis, and injury in *Aldh2*-KO mice. (**A**) Representative H&E staining of formalin-fixed liver sections from WT and *Aldh2*-KO mice exposed to a single dose of ethanol 3.5, 4.0, or 5.0 g/kg (n ≥ 3~4/group). The arrows indicate the area of inflammation. (**B**) Hepatic IL-6, TNF-α, and lipid peroxide levels are presented for the indicated groups. (**C**) Representative TUNEL staining for apoptotic hepatocytes (arrows) in WT and *Aldh2*-KO mice exposed to a single dose of ethanol. (**D**) Hepatic caspase-3 activity and serum AST and ALT levels are shown for the indicated groups. CON: Control, E3.5: EtOH 3.5 g/kg, E4: EtOH 4.0 g/kg, E5: EtOH 5.0 g/kg. Bar scale, 100 μm in A and B. Data show means ± SD where statistical significance was determined using two-way ANOVA; * p<0.05, ** p<0.01, *** p<0.001, **** p<0.0001.

### 3.3 Binge alcohol exposure increased oxidative stress and apoptotic markers with decreased intestinal TJ & AJ proteins in Aldh2-KO mice

Our results thus far showed that *Aldh2*-KO mice were more sensitive to binge alcohol-induced gut and liver injuries than the corresponding WT mice. Therefore, we focused on studying the mechanisms underlying elevated gut injury and endotoxemia in *Aldh2*-KO mice. We hypothesized that the levels of oxidative stress, enterocyte apoptosis, and proteasomal degradation of intestinal TJ/AJ proteins, which are important in the maintenance of integrity and function of the intestinal barrier, are changed after exposure to a single dose of binge alcohol in *Aldh2*-KO mice. With this hypothesis, the intestinal homogenates were analyzed to determine the immunoreactive levels of gut TJ/AJ proteins. Immunoblot results showed that a single dose of ethanol 3.5, 4.0, or 5.0 g/kg, p.o. significantly increased levels of intestinal nitroxidative stress markers, including CYP2E1 and iNOS, and apoptotic indicator proteins, such as Bax and cleaved (active)-caspase 3 in binge alcohol-exposed *Aldh2*-KO mice 1 h after alcohol exposure. These data are consistent with the results of histological, serum, and caspase-3 activity analyses (Fig. 1). However, little or no changes were observed in intestinal proteins of the corresponding WT mice (Fig. 3A and B). In addition, the expressions of gut TJ proteins, including Zonula occluden-1 (ZO-1), Occludin, Claudin-1, Claudin-3, and Claudin-4, and gut AJ proteins, such as β-catenin and E-cadherin, and Plakoglobin were significantly decreased in alcohol-exposed *Aldh2*-KO mice. In contrast, little changes in the gut TJ/AJ proteins were observed in the corresponding WT mice (Fig. 3C and D). It is possible that significant differences in the abundance and composition of gut microbiota in the basal conditions of WT and *Aldh2*-KO mice could play a role in this differential sensitivity toward alcohol-related changes, as discussed [11, 12]. Bacterial metagenomic sequencing analyses showed that the abundance of beneficial bacteria promoting gut health (e.g., Verrucomicrobia) were decreased, while pathogenic bacteria (e.g., Proteobacteria) were increased in *Aldh2*-KO mice compared to those of WT mice (Fig. 3E). All these results indicated that increased nitroxidative stress and enterocyte apoptosis, decreased gut TJ/AJ protein levels, and differential abundance or composition of gut microbiota could lead to elevated gut leakiness in binge alcohol-exposed *Aldh2*-KO mice compared with the corresponding WT mice. Our findings are in agreement with the greater sensitivity of *Aldh2*-KO mice exposed to an ethanol liquid diet for 4 weeks [42] and further suggest an important role of ALDH2 in protecting against binge alcohol-mediated gut leakiness.

**Figure 3:**
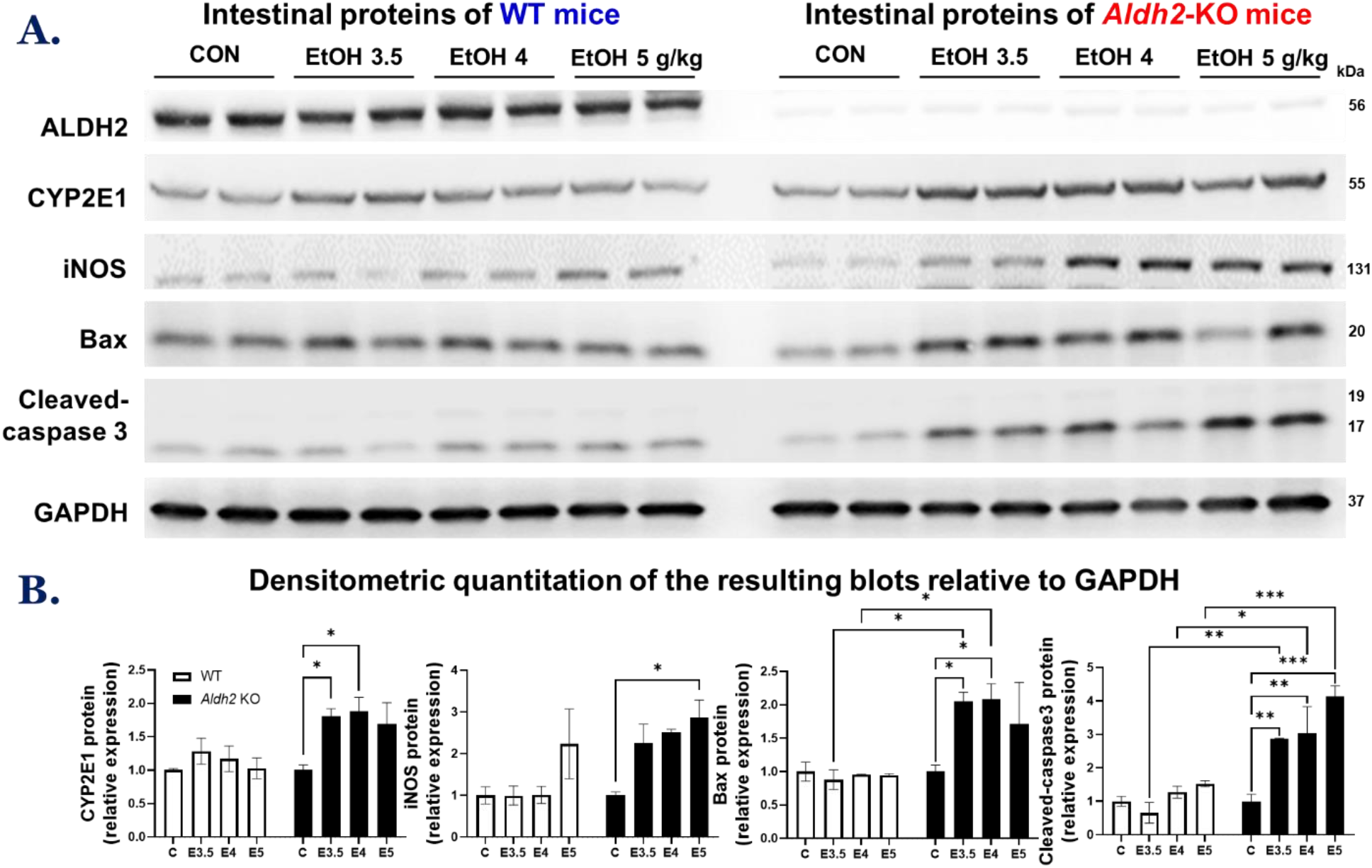
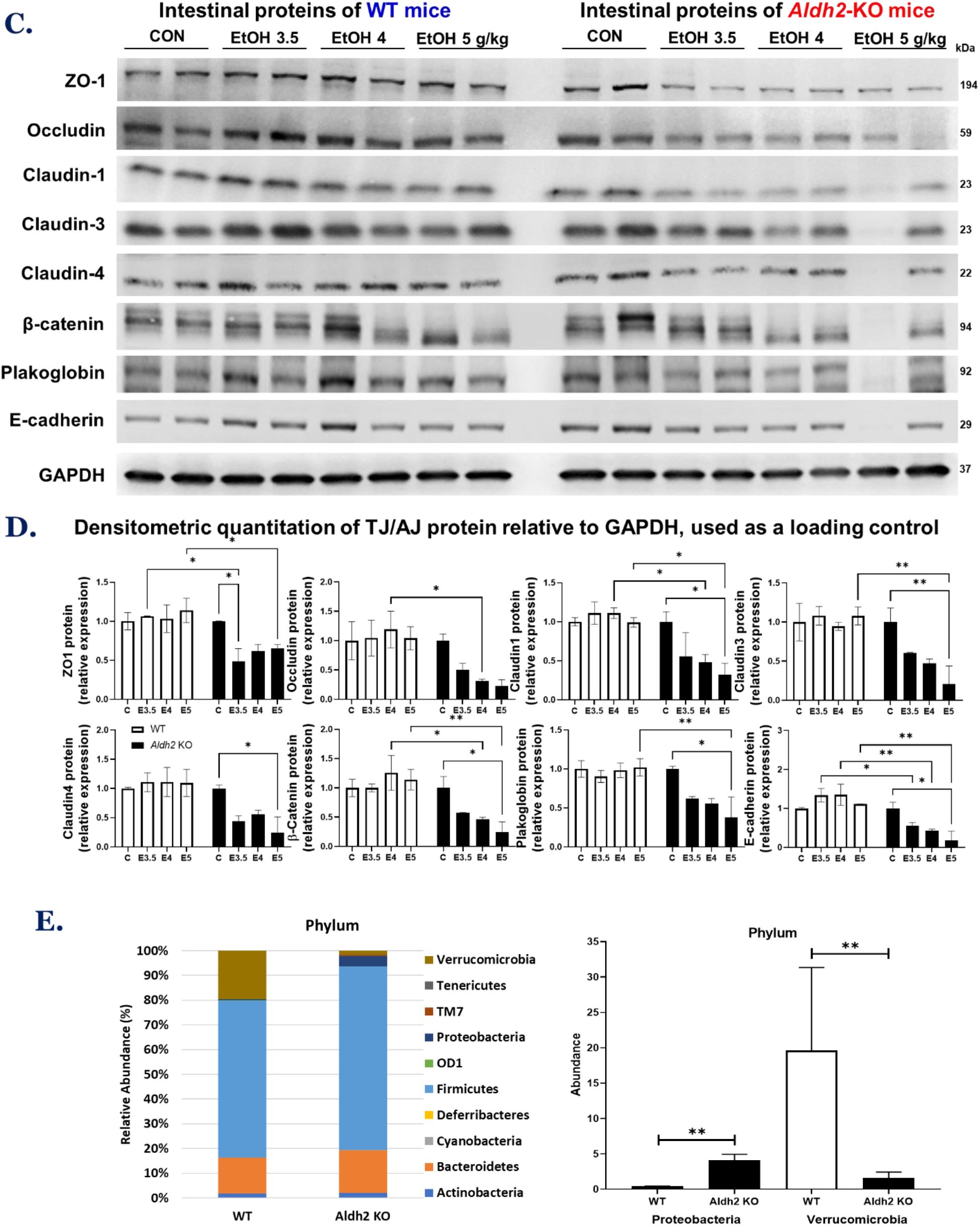
The absence of ALDH2 and differential composition of gut microbiota contributed to increased oxidative stress and apoptotic markers with decreased gut TJ & AJ proteins in alcohol-exposed *Aldh2*-KO mice but not in the WT mice. (**A**) Representative immunoblot analysis of intestinal ALDH2, CYP2E1, iNOS, Bax, Cleaved (active)-caspase 3, and a loading control GAPDH for the indicated mouse groups (n≥3~4/group). Each lane represents a mixture of equal amounts of protein from two different samples within the same group. (**B**) Densitometric quantitation of the resulting blots relative to GAPDH is presented. (**C**) Representative immunoblot analysis of gut TJ/AJ proteins: ZO-1, Occludin, Claudin-1, Claudin-3, Claudin-4, β-catenin, E-cadherin, and Plakoglobin for the indicated mouse groups. (**D**) Densitometric quantitation of TJ/AJ protein relative to GAPDH is shown. (**E**) Comparison of the taxonomic abundance bacteria phyla in the basal conditions of WT and *Aldh2*-KO mice (n≥3/group) is shown. CON: Control, E3.5: EtOH 3.5 g/kg, E4: EtOH 4.0 g/kg, E5: EtOH 5.0 g/kg. Data are expressed as means ± SD where statistical significance was determined using two-way ANOVA: * p<0.05, ** p<0.01, *** p<0.001, **** p<0.0001.

### 3.4 Alcohol exposure increased nitration of gut TJ/AJ proteins, leading to their decreased levels and gut leakiness in Aldh2-KO mice

After finding the decreases in gut TJ/AJ proteins in binge alcohol-exposed *Aldh2*-KO mice, we further studied the underlying mechanism for their decrements. Recent studies showed that nitration of gut TJ/AJ proteins leads to their ubiquitin-dependent proteasomal degradation, contributing to elevated gut leakiness, endotoxemia, and liver injury [9, 10, 43]. Here, immunoblot results showed that the levels of nitrated (3-nitrotyrosine or 3-NT) and ubiquitinated proteins in the gut extracts of *Aldh2*-KO mice were significantly increased after 1-h exposure to a single dose of ethanol 3.5, 4.0, or 5.0 g/kg p.o. (Fig. 4A). In contrast, the levels of protein nitration and ubiquitination in the control and alcohol-exposed WT mice were similar or slightly decreased in the latter groups. Furthermore, immunoprecipitation with anti-Occludin, anti-Claudin1 (TJ proteins), or anti-β-catenin (AJ protein) followed by immunoblot analysis with the specific antibody to each indicated target protein revealed that these intestinal TJ/AJ proteins were nitrated despite being decreased in binge alcohol-exposed *Aldh2*-KO mice (Fig. 4B). In contrast, little changes were observed in the corresponding WT mice. These findings suggest that nitration of gut TJ/AJ proteins, as shown with the selected proteins, could contribute to their proteolytic degradation, as recently reported [9, 10, 43], gut leakiness, endotoxemia, and ultimately hepatic injury in binge alcohol-exposed *Aldh2*-KO mice through the gut-liver axis.

**Figure 4:**
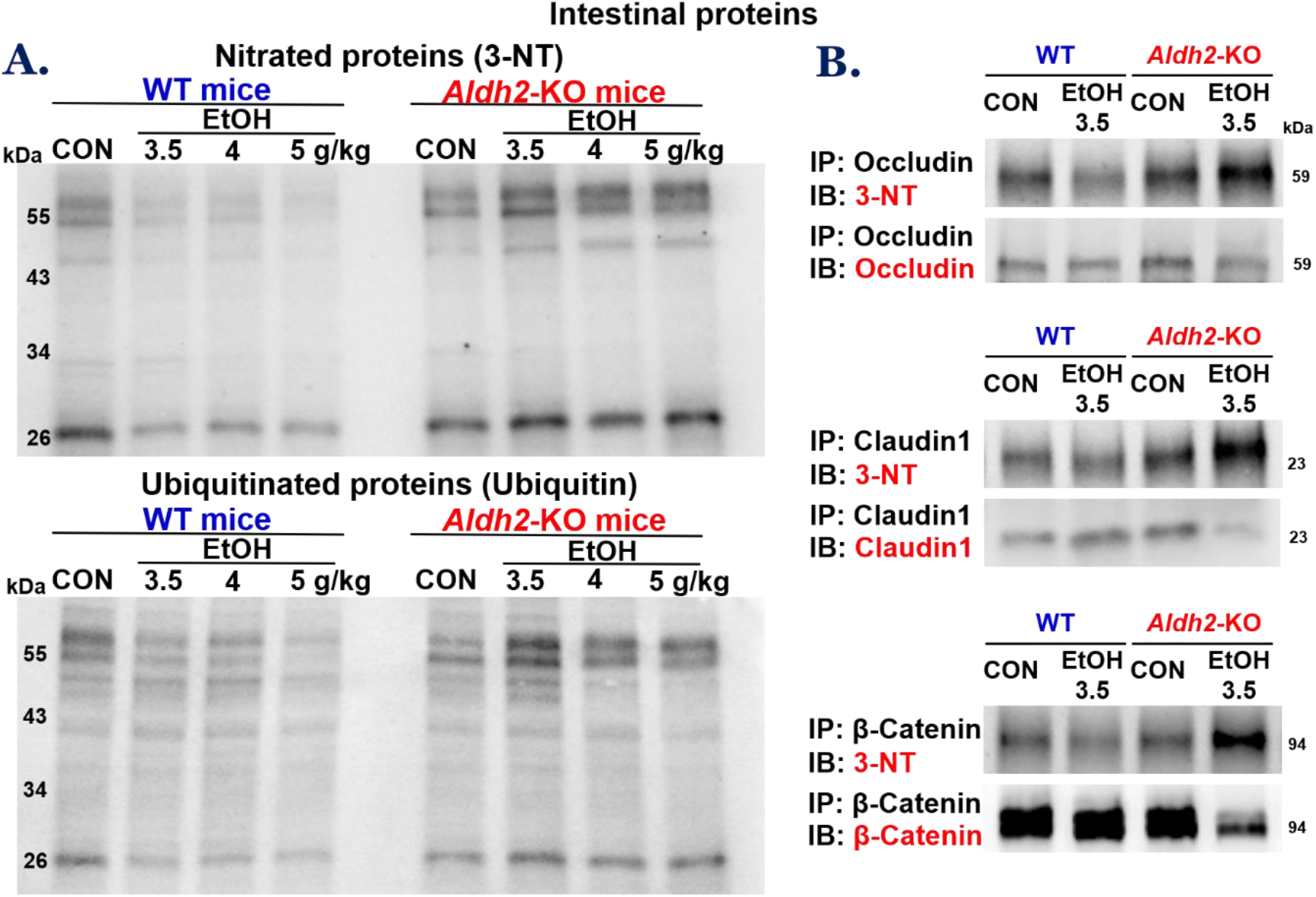
Binge alcohol increased nitration of gut TJ/AJ proteins, leading to their decreased levels in *Aldh2*-KO mice. (**A**) Representative immunoblot analysis of nitrated and ubiquitinated proteins detected by the specific antibody against 3-NT and Ubiquitin. (**B**) Equal amounts (0.5 mg/sample) of intestinal proteins from the control or ethanol-exposed WT or *Aldh2*-KO mice were immunoprecipitated with the specific antibody to each indicated TJ or AJ protein and then subjected to immunoblot analysis with the specific antibody to 3-NT (top panels) or each TJ or AJ protein of interest (lower panels).

### 3.5 Suppression of ALDH2 promoted epithelial cell permeability with decreased TJ & AJ protein levels, while ALDH2 activation ameliorated these changes in alcohol-exposed T84 colonic cells

To confirm the results with *Aldh2*-KO mice and further elucidate the underlying mechanism by which ALDH2 protects against alcohol-induced gut leakiness (and liver injury), we utilized T84 human colonic cells by evaluating the changes in epithelial cell permeability and TJ/AJ protein levels after ethanol exposure. Based on high alcohol concentrations present in the intestine [36], T84 colonic cells were treated with ethanol (0.15 M) in the absence or presence of Daidzin, an ALDH2 inhibitor [37, 38], or Alda-1, an ALDH2 activator [39–41]. Fig. 5A represents the results of ALDH2 protein expression determined by immunoblot analysis and its activity in alcohol-exposed T84 cells with or without an ALDH2 inhibitor (Daidzin) or an ALDH2 activator (Alda-1). The levels of ALDH2 protein and activity were slightly decreased by alcohol exposure and significantly suppressed by Daidzin treatment. However, Alda-1 restored the levels of ALDH2 protein and activity, consistent with the previous findings [38]. The results of epithelial cell permeability monitored with FITC-D4 fluorescence signals showed that little or no changes were observed with Daidzin or Alda-1 in the absence of alcohol. However, in the presence of alcohol, ALDH2 inhibition significantly promoted alcohol-induced cell permeability, while ALDH2 activation significantly blocked this event (Fig. 5B). Consistently, alcohol exposure decreased T84 cell viability, and this was significantly reduced in the presence of Daidzin but prevented by Alda1 pretreatment (Fig. 5C). In addition, decreased TJ/AJ protein levels after alcohol exposure were exacerbated by ALDH2 inhibition, while these changes were significantly attenuated by ALDH2 activation (Fig. 5D). These findings further support the protective role of ALDH2 in alcohol-mediated epithelial barrier permeability in T84 cells, consistent with the gut leakiness observed in the *Aldh2*-KO mice.

**Figure 5:**
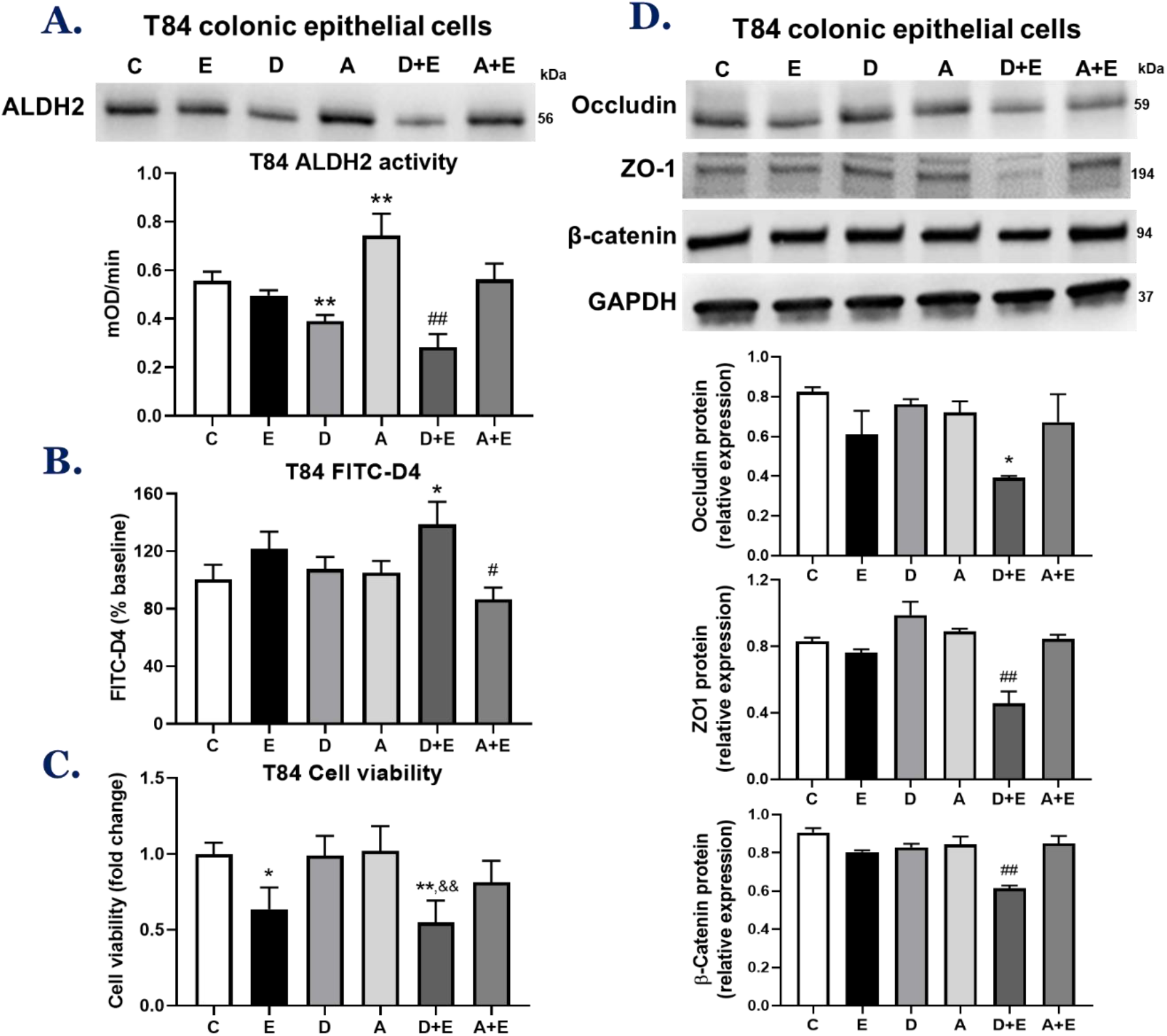
ALDH2 inhibition promoted alcohol-mediated T84 human colonic cell permeability with decreased TJ/AJ proteins, while all these changes were prevented by ALDH2 activation. Representative levels of (**A**) ALDH2 protein expression and activity, (**B**) epithelial cell permeability to FITC-D4, (**C**) cell viability, and (**D**) TJ/AJ proteins in T84 colonic epithelial cells exposed to 150 mM ethanol in the absence or presence of Daidzin (an ALDH2 inhibitor, 10 μM) or Alda-1 (an ALDH2 activator, 20 μM). C: Control, E: EtOH, D: Daidzin, A: Alda-1, D+E: Daidzin+EtOH, A+D: Alda-1+EtOH. Data indicate means ± SD of triplicate wells from two separate experiments. * p < 0.05, ** p < 0.01, *** p < 0.001 vs. Control. & p < 0.05 vs. Daidzin. # p < 0.05, ## p < 0.01, ### p < 0.001 vs. EtOH.

### 3.6 Acute alcohol exposure disrupted TJ & AJ organization and expression in T84 colonic cells in an ALDH2-dependent manner

In addition to the measurements of ALDH2 levels, cell permeability, and TJ/AJ proteins, confocal immunofluorescence microscopy was utilized to further visualize TJ/AJ organization and expression in cultured T84 colonic cells. The cells were exposed to 130 mM ethanol with or without an ALDH2 inhibitor (Daidzin) or an ALDH2 activator (Alda-1), and immunofluorescence was determined for the levels of TJ proteins (Occludin and ZO1) and AJ protein (β-catenin). Confocal analysis results showed that acute alcohol exposure slightly changed the levels of ALDH2 in T84 colonic cells. ALDH2 expression was markedly decreased after treatment with an ALDH2 inhibitor (Daidzin) but was significantly increased in the presence of an ALDH2 activator (Alda-1) (Fig. 6A), consistent with the previous report [38]. Furthermore, ethanol exposure alone slightly modified TJ/AJ structure and distribution, while ALDH2 inhibition exacerbated the effects of ethanol on disrupted TJ/AJ organization and expression in T84 colonic cells. On the other hand, ALDH2 activation by Alda-1 prevented ethanol-induced deformation and decrement of TJ/AJ structure and expression (Fig. 6B-D), supporting the important role of ALDH2 in protecting against alcohol-induced epithelial cell barrier dysfunction or gut permeability.

**Figure 6:**
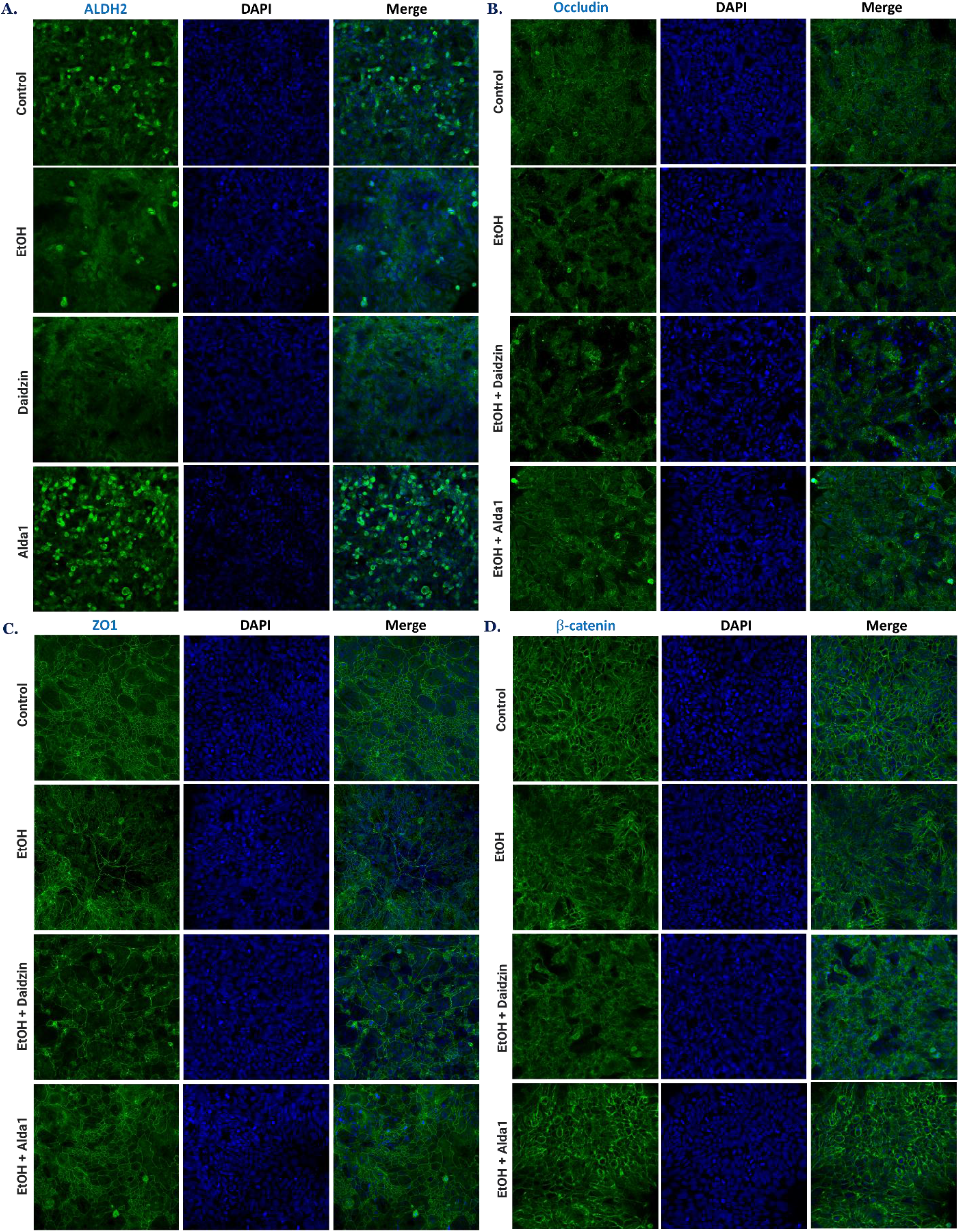
Confocal images revealed that acute alcohol exposure disrupted TJ/AJ organization and expression in T84 colonic cells in an ALDH2-dependent manner. Confocal images show acute alcohol exposure negatively affected the levels of ALDH and TJ/AJ proteins. Representative confocal images for (**A**) ALDH2, (**B**) Occludin, (**C**) ZO1, and (**D**) β-catenin without or with exposure to 130 mM ethanol [36] in the absence or presence of an ALDH2 inhibitor (Daidzin, 10 μM) or an ALDH2 activator (Alda-1, 20 μM) [38]. The cells were imaged at 200x magnification, and cell nuclei were counterstained with DAPI.

**Figure 7:**
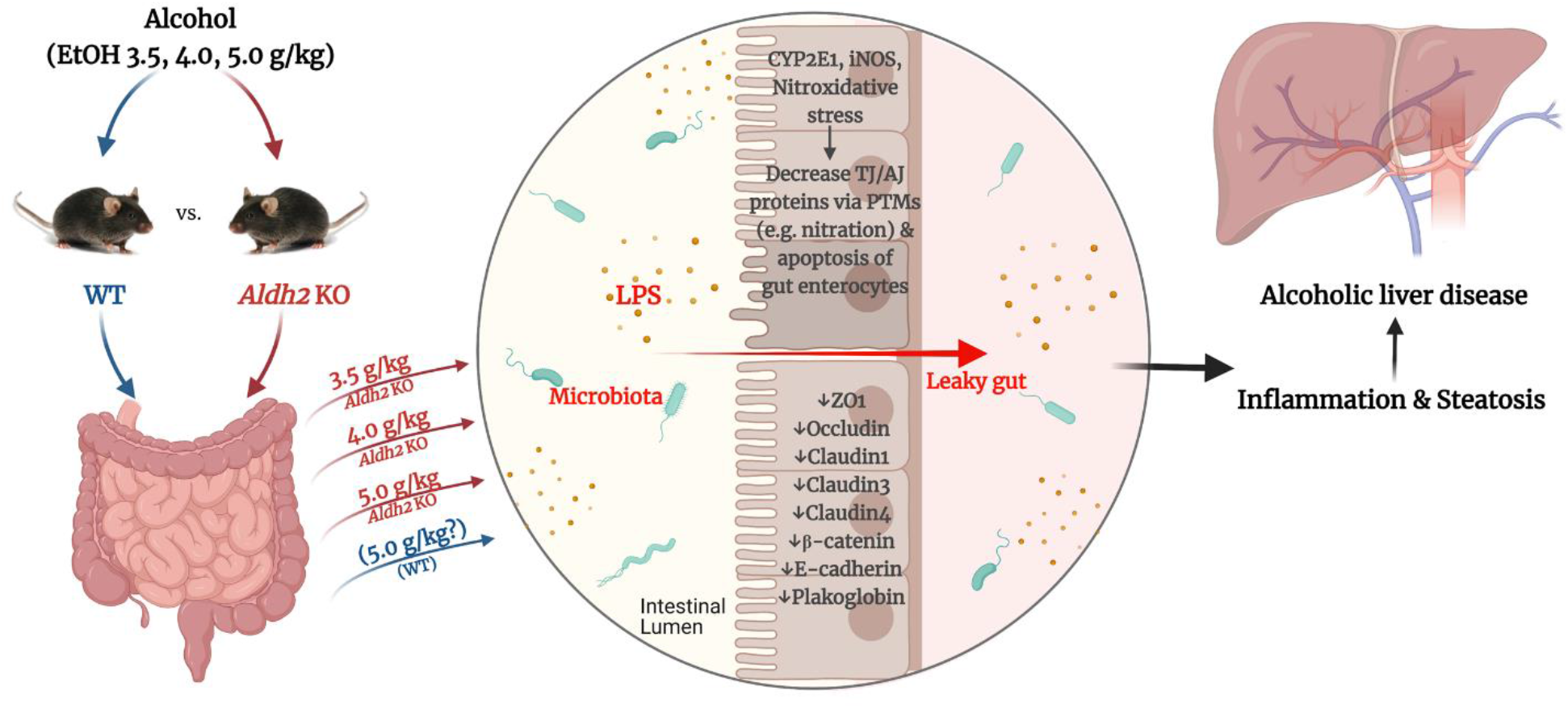
*Aldh2*-knockout mice are more susceptible to binge alcohol-induced gut leakiness, endotoxemia, and acute liver injury than WT mice. The deletion of *Aldh2* gene in *Aldh2*-KO mice contributes to greater production of oxidative and nitrative stress markers, which result in elevated PTMs, such as nitration, followed by ubiquitin-dependent proteasomal degradation of gut TJ/AJ proteins, apoptosis of enterocytes, and intestinal barrier dysfunction, ultimately leading to gut leakiness and endotoxemia after exposure to a single dose of ethanol even at 3.5 g/kg. In contrast, no or little changes were observed in the corresponding WT mice. Consequently, elevated endotoxemia and/or bacterial translocation can cause inflammation, oxidative stress, apoptosis, and subsequent hepatic steatosis, and acute liver injury through the gut-liver axis.

## 4. Discussion

Excessive or binge alcohol consumption has been an important risk factor in promoting various diseases, injuries, and pathological conditions worldwide. Alcohol (ethanol) can penetrate virtually all tissues and negatively affect their functions especially after excessive and chronic intake [2, 13, 44, 45]. In fact, alcohol-associated tissue injury and hospitalization represent a significant portion (approximately a quarter) of all hospitalizations although the majority of all these incidences can be prevented by drinking in moderation or simple abstinence [46, 47]. These alcohol-related toxicities can be observed in people with AUD, animal models (rodents and primates), and *in vitro* culture cells, suggesting conservative toxic mechanisms among different species or models. There are multiple mechanisms by which excessive and binge alcohol can stimulate tissue or cell injury and worsen disease states such as liver fibrosis/cirrhosis and cancer [48]. Two of the main causal factors for tissue injury could be increased redox changes through oxidative ethanol metabolism and elevated nitroxidative stress after alcohol exposure. The sources of elevated nitroxidative stress include activation and/or upregulation of pro-oxidant proteins (such as CYP2E1, NADPH oxidase, xanthine oxidase, and iNOS), dysregulation of mitochondrial electron transport proteins, reduced levels of cellular NAD^+^/NADH ratio and an antioxidant glutathione, accompanied by suppression of the antioxidant enzymes (e.g., superoxide dismutase, catalase, glutathione peroxidase, peroxiredoxin, and ALDH isozymes) [49–51].

It is well-established that genetic risk factors or mutations also play an important role in alcohol-associated tissue injury and diseases. For instance, mutation of *ALDH2* gene (*ALDH2*2*) in humans and *Aldh2* gene deletion in mice (e.g., *Aldh2*-KO mice) are known to have very low or no ALDH2 activity with increased sensitivity to alcohol-induced tissue injury and cancer compared to people with normal *ALDH2* gene and WT mice, respectively [19–21, 23–27, 52]. In humans, the suppression of a defensive enzyme ALDH2 leads to an accumulation of abnormal acetaldehyde-protein and/or -DNA adducts observed in people with *ALDH2* gene mutation after alcohol intake [52–54]. In mice, the deletion of ALDH2 also contributes to increased acetaldehyde-DNA adducts or damages, cardiovascular abnormalities, inflammatory liver diseases, and even short lifespan or cancer [26, 27, 55–57] after long-term alcohol exposure. In addition, the deletion of *Aldh2* gene leads to higher acetaldehyde accumulation in the liver and brain in mice after exposure to a single oral dose of alcohol [25]. However, the effects of binge alcohol on *Aldh2*-KO mice or people with *ALDH2*2* gene have not been studied especially on the gut-liver axis, although elevated acetaldehyde did not seem to play a critical role in binge alcohol-induced acute liver injury [58]. In this study, we found that *Aldh2*-KO mice are more susceptible to binge alcohol-mediated gut and inflammatory liver injury with increased oxidative stress and apoptosis of enterocytes and hepatocytes compared with WT counterparts after exposure to a single dose of ethanol even at 3.5 g/kg, p.o. (Figs. 1 and 2). These changes were accompanied by intestinal barrier dysfunction with elevated serum endotoxin (LPS), which can stimulate inflammatory tissue injury [9, 10, 43]. Additionally, based on the significant induction of CYP2E1 (Fig. 3C) [59, 60] and its properties in producing ROS [17, 61] and promoting various PTMs like nitration [9, 10, 43], elevated CYP2E1, at least partially, plays a contributing role in binge-alcohol mediated gut leakiness and acute liver injury in *Aldh2*-KO mice. These results are in agreement with recent reports, where *Cyp2e1*-KO mice were protected from binge alcohol-mediated gut leakiness and acute liver injury [9, 29, 62].

More recent reports have indicated the important roles of gut leakiness and gut-derived bacteria with their products like LPS, which can be transported to other tissues such as the liver and brain to cause inflammatory tissue injury through the gut-liver-brain axis [63–65]. The manifestation of gut leakiness and endotoxemia has been implicated by not only alcohol but also other diets (i.e., fructose and Western-style high-fat diets, HFDs) [10, 66, 67] or in different pathological conditions e.g., neurodegenerative diseases, HIV-mediated full-blown AIDS symptoms, obesity, and nonalcoholic steatohepatitis (NASH) [65, 68, 69]. Despite these active mechanistic studies, the exact molecular mechanisms of gut leakiness and its impacts on promoting inflammatory injury in other tissues are still incompletely understood. To our best knowledge, gut dysbiosis with a decreased abundance of beneficial bacteria and an increased population of pathobionts could play a contributing role in elevated gut leakiness and local inflammation, contributing to endotoxemia, as reported [11, 12], [70]. In fact, our metagenomic sequencing results (Fig. 3E) showed that the basal conditions of gut microbiota abundance and composition in *Aldh*2-KO mice were distinct from those of WT mice. For instance, increased pathogenic Proteobacteria with decreased Verrucomicrobia in *Aldh*2-KO mice (Fig. 3E) suggested that these mice would be more sensitive to cell/tissue injury following exposure to risk factors such as binge alcohol, as shown in this study. Based on these differences, it is expected that the levels of bacterial metabolites short-chain fatty acids (SCFAs) would also be imbalanced in *Aldh2*-KO mice, although this analysis warrants future characterization. Additionally, owing to the role of ALDH2 in the metabolism of lipid aldehydes (4-HNE and MDA) [71, 72], people with *ALDH2*2* mutation and *Aldh2*-KO mice could also be more susceptible to lipid peroxide-DNA adducts and tissue injury upon exposure to other nonalcoholic diets such as fructose and Western-style HFDs.

In the gut, the intestinal mucosal barrier is composed of different TJ/AJ complex proteins essential for maintaining the integrity and function of the gut barrier. Tight junctions are located at the apical end of epithelial cells (e.g., Occludin, ZO1, and Claudins), while adherent junctions (such as E-cadherin and α/β-Catenins) reside under the tight junctions and assist in supporting TJ integrity through a complex interaction with desmosomes (i.e., Plakoglobin) and cytoskeletons [73]. Therefore, disruption of these TJ/AJ proteins subsequently leads to gut leakiness, allowing bacteria and their products such as endotoxin (LPS) to enter the circulation and induce liver inflammation and other organ damages [9, 10, 29]. Rao and his co-workers consistently reported a causal role of acetaldehyde in the permeability of colonic epithelial Caco-2 cells [74] and gut leakiness in *Aldh2*-KO mice exposed to the Lieber-DeCarli alcohol liquid diet for 4 weeks [42] although the mechanisms for decreased gut TJ/AJ proteins by PTMs, including nitration, was not investigated. In the current study with acute alcohol exposure, we observed consistent elevations of epithelial permeability (and apoptosis) in T84 colonic cells and gut leakiness in *Aldh2*-KO mice compared with the untreated controls and corresponding WT mice. Markedly increased enterocyte apoptosis was also observed in *Aldh2*-KO compared to the controls or WT mice exposed to the same doses of ethanol. Interestingly, our findings clearly showed the elevated gut protein nitration and ubiquitination with decreased gut TJ/AJ levels in alcohol-exposed *Aldh2*-KO mice, while these changes were not observed in WT counterparts. Although the protein nitration was performed with a few selected TJ/AJ proteins, the current results are in agreement with previous reports [9, 10, 43], where the other gut TJ/AJ proteins tended to undergo similar types of nitration or other PTMs like phosphorylation and acetylation, contributing to the decreased TJ/AJ proteins through ubiquitin-dependent proteasomal degradation [9, 10, 16, 29]. In addition, increased gut leakiness and paracellular permeability observed in alcohol-exposed *Aldh2*-KO mice and Daidzin-exposed T84 cells, respectively, could be mediated by accumulated acetaldehyde, which was shown to disrupt the interaction between various TJ/AJ proteins, as previously described [74]. Nonetheless, our findings suggest that acute alcohol exposure increases nitroxidative stress, which leads to elevated apoptosis of enterocytes and nitration of gut TJ/AJ proteins followed by their degradation, resulting in gut leakiness, endotoxemia, and subsequently inflammatory liver injury in *Aldh2*-KO mice.

Consistent with the *in vivo* results, acute alcohol exposure caused changes in TJ/AJ protein expression, ALDH2 protein levels (similar to a previous report [75]) and activity, and cell permeability in T84 colonic cells. ALDH2 inhibition by an ALDH2 inhibitor (Daidzin) further worsened the alcohol-induced epithelial permeability and reduced TJ/AJ proteins than alcohol alone in T84 cells. In contrast, activation of ALDH2 by its activator (Alda-1) protected the T84 cells from the effects of alcohol, and these results are consistent with previous reports of Alda-1-mediated gut protection [76, 77]. Collectively, our findings suggest the protective role of ALDH2 in alcohol-mediated nitroxidative stress, nitration and decrement of gut TJ/AJ proteins, gut leakiness, endotoxemia, and acute liver injury through the gut-liver axis. In the line of our study, previous research on alcohol-associated tissue injury in *Aldh2*-KO mice has also revealed an important role of acetaldehyde and ALDH2 in the intestinal barrier and liver function [42]. The authors reported that *Aldh2*-KO mice were more sensitive to alcohol-induced gut injury through redistribution of intestinal TJ/AJ proteins as well as increases in serum ALT and AST and fat accumulation in the liver, where both alcohol and elevated acetaldehyde likely resulting from the *Aldh2* deletion played a role in gut and liver damage. Despite the different treatments between the earlier study for chronic administration [42] and the current study of acute exposure, the outcome would be similar to show the susceptibility of *Aldh2*-KO mice to alcohol-mediated gut leakiness and tissue injury through the gut-liver axis. Furthermore, our current results showed, for the first time, the critical role of nitroxidative stress to nitrate intestinal TJ/AJ proteins, leading to their degradation, elevated endotoxemia, and acute liver injury in *Aldh2*-KO mice upon exposure to binge alcohol. Since ALDH2 is also responsible for the metabolism of toxic lipid aldehydes, it is likely that *Aldh2*-KO mice could also be more sensitive to nonalcoholic substance-mediated tissue damage, suggesting ALDH2 as an important target of clinical preventive or therapeutic importance.

## 5. Conclusion

This study demonstrates the direct effect of *Aldh2* gene deletion (or inactivation) on binge alcohol-mediated gut leakiness and acute liver injury upon exposure to a single oral dose of alcohol even at 3.5 or 4.0 g/kg. The genetic deletion or inactivation of ALDH2 contributes to greater susceptibility to alcohol-induced elevated oxidative and nitrative stress and various PTMs including nitration, leading to gut TJ/AJ protein degradation and enterocyte apoptosis. These changes contribute to gut leakiness, endotoxemia, and acute liver injury through the gut-liver axis in *Aldh2*-KO mice compared with the corresponding WT mice. Based on these findings, ALDH2 is an important target for preventing alcohol-mediated gut leakiness and inflammatory liver injury.

## Abbreviations

AUD: alcohol used disorder
ALD: alcoholic liver disease
ALDH2: mitochondrial aldehyde dehydrogenase 2
CYP2E1: ethanol-inducible cytochrome P450-2E1
ECL: enhanced chemiluminescence
FITC-D4: fluorescein isothiocyanate dextran-4 kDa
MDA: malondialdehyde
4-HNE: 4-hydroxynonenal
H&: Ehematoxylin & eosin
LPS: lipopolysaccharide
TJ: tight junction proteins
AJ: adherent junction proteins
TUNEL: terminal deoxynucleotidyl transferase dUTP nick end labeling
PTM: post-translational modification

## Acknowledgments

This research was supported by the Intramural Program Research Fund of the National Institute of Alcohol Abuse and Alcoholism, National Institutes of Health.

## Author contributions

Conceptualization: B.J.S, W.R.; original draft preparation: W.R.; animal acquisition: T.K.; experiment and data analysis: W.R.; manuscript revision for important intellectual content: B.J.S., X.W., T.K., S.B.C.; supervision: B.J.S.

## References

1. Estaki, M., et al., QIIME 2 Enables Comprehensive End-to-End Analysis of Diverse Microbiome Data and Comparative Studies with Publicly Available Data. Curr Protoc Bioinformatics, 2020. 70(1): p. e100.

2. Wang, H.J., S. Zakhari, and M.K. Jung, Alcohol, inflammation, and gut-liver-brain interactions in tissue damage and disease development. World J Gastroenterol, 2010. 16(11): p. 1304–13.

3. Schäfer, C., et al., Concentrations of lipopolysaccharide-binding protein, bactericidal/permeability-increasing protein, soluble CD14 and plasma lipids in relation to endotoxaemia in patients with alcoholic liver disease. Alcohol Alcohol, 2002. 37(1): p. 81–6.

4. Marchitti, S.A., et al., Non-P450 aldehyde oxidizing enzymes: the aldehyde dehydrogenase superfamily. Expert Opin Drug Metab Toxicol, 2008. 4(6): p. 697–720.

5. Mutlu, E., et al., Intestinal dysbiosis: a possible mechanism of alcohol-induced endotoxemia and alcoholic steatohepatitis in rats. Alcohol Clin Exp Res, 2009. 33(10): p. 1836–46.

6. Song, B.J., et al., Post-translational modifications of mitochondrial aldehyde dehydrogenase and biomedical implications. J Proteomics, 2011. 74(12): p. 2691–702.

7. Jackson, B.C., et al., Dead enzymes in the aldehyde dehydrogenase gene family: role in drug metabolism and toxicology. Expert Opin Drug Metab Toxicol, 2015. 11(12): p. 1839–47.

8. Liangpunsakul, S., et al., Quantity of alcohol drinking positively correlates with serum levels of endotoxin and markers of monocyte activation. Sci Rep, 2017. 7(1): p. 4462.

9. Cho, Y.E., et al., Apoptosis of enterocytes and nitration of junctional complex proteins promote alcohol-induced gut leakiness and liver injury. J Hepatol, 2018. 69(1): p. 142–153.

10. Cho, Y.E., et al., Fructose Promotes Leaky Gut, Endotoxemia, and Liver Fibrosis Through Ethanol-Inducible Cytochrome P450-2E1-Mediated Oxidative and Nitrative Stress. Hepatology, 2021. 73(6): p. 2180–2195.

11. Bishehsari, F., et al., Alcohol and Gut-Derived Inflammation. Alcohol research: current reviews, 2017. 38(2): p. 163–171.

12. Ballway, J.W. and B.J. Song, Translational Approaches with Antioxidant Phytochemicals against Alcohol-Mediated Oxidative Stress, Gut Dysbiosis, Intestinal Barrier Dysfunction, and Fatty Liver Disease. Antioxidants (Basel), 2021. 10(3).

13. Bode, C., V. Kugler, and J.C. Bode, Endotoxemia in patients with alcoholic and non-alcoholic cirrhosis and in subjects with no evidence of chronic liver disease following acute alcohol excess. Journal of Hepatology, 1987. 4(1): p. 8–14.

14. Xiao, J., et al., Lychee (Litchi chinensis Sonn.) Pulp Phenolic Extract Provides Protection against Alcoholic Liver Injury in Mice by Alleviating Intestinal Microbiota Dysbiosis, Intestinal Barrier Dysfunction, and Liver Inflammation. J Agric Food Chem, 2017. 65(44): p. 9675–9684.

15. Cederbaum, A.I., Alcohol metabolism. Clinics in liver disease, 2012. 16(4): p. 667–685.

16. Song, B.J., et al., Mitochondrial dysfunction and tissue injury by alcohol, high fat, nonalcoholic substances and pathological conditions through post-translational protein modifications. Redox Biol, 2014. 3: p. 109–23.

17. Song, B.J., et al., Translational Implications of the Alcohol-Metabolizing Enzymes, Including Cytochrome P450-2E1, in Alcoholic and Nonalcoholic Liver Disease. Adv Pharmacol, 2015. 74: p. 303–72.

18. Rungratanawanich, W., et al., Advanced glycation end products (AGEs) and other adducts in aging-related diseases and alcohol-mediated tissue injury. Exp Mol Med, 2021. 53(2): p. 168–188.

19. Gill, K., et al., An examination of ALDH2 genotypes, alcohol metabolism and the flushing response in Native Americans. J Stud Alcohol, 1999. 60(2): p. 149–58.

20. Wall, T.L., et al., Hangover symptoms in Asian Americans with variations in the aldehyde dehydrogenase (ALDH2) gene. J Stud Alcohol, 2000. 61(1): p. 13–7.

21. Yokoyama, T., et al., Alcohol flushing, alcohol and aldehyde dehydrogenase genotypes, and risk for esophageal squamous cell carcinoma in Japanese men. Cancer Epidemiol Biomarkers Prev, 2003. 12(11): p. 1227–33.

22. Chen, C.H., et al., Targeting aldehyde dehydrogenase 2: new therapeutic opportunities. Physiol Rev, 2014. 94(1): p. 1–34.

23. Chen, C.H., et al., Novel and prevalent non-East Asian ALDH2 variants; Implications for global susceptibility to aldehydes’ toxicity. EBioMedicine, 2020. 55: p. 102753.

24. Kitagawa, K., et al., Aldehyde dehydrogenase (ALDH) 2 associates with oxidation of methoxyacetaldehyde; in vitro analysis with liver subcellular fraction derived from human and Aldh2 gene targeting mouse. FEBS Lett, 2000. 476(3): p. 306–11.

25. Isse, T., et al., Aldehyde dehydrogenase 2 gene targeting mouse lacking enzyme activity shows high acetaldehyde level in blood, brain, and liver after ethanol gavages. Alcohol Clin Exp Res, 2005. 29(11): p. 1959–64.

26. Ma, H., et al., Aldehyde dehydrogenase 2 knockout accentuates ethanol-induced cardiac depression: role of protein phosphatases. J Mol Cell Cardiol, 2010. 49(2): p. 322–9.

27. Matsumoto, A., et al., Ethanol reduces lifespan, body weight, and serum alanine aminotransferase level of aldehyde dehydrogenase 2 knockout mouse. Alcohol Clin Exp Res, 2014. 38(7): p. 1883–93.

28. Kwon, H.J., et al., Aldehyde dehydrogenase 2 deficiency ameliorates alcoholic fatty liver but worsens liver inflammation and fibrosis in mice. Hepatology, 2014. 60(1): p. 146–57.

29. Abdelmegeed, M.A., et al., CYP2E1 potentiates binge alcohol-induced gut leakiness, steatohepatitis, and apoptosis. Free Radic Biol Med, 2013. 65: p. 1238–1245.

30. Lambert, J.C., et al., Prevention of alterations in intestinal permeability is involved in zinc inhibition of acute ethanol-induced liver damage in mice. J Pharmacol Exp Ther, 2003. 305(3): p. 880–6.

31. Rhee, S.G., et al., Methods for detection and measurement of hydrogen peroxide inside and outside of cells. Mol Cells, 2010. 29(6): p. 539–49.

32. Kalyanaraman, B., et al., Measuring reactive oxygen and nitrogen species with fluorescent probes: challenges and limitations. Free Radic Biol Med, 2012. 52(1): p. 1–6.

33. Forman, H.J., et al., Even free radicals should follow some rules: a guide to free radical research terminology and methodology. Free Radic Biol Med, 2015. 78: p. 233–5.

34. Tran, M., et al., A Potential Role for SerpinA3N in Acetaminophen-Induced Hepatotoxicity. Mol Pharmacol, 2021. 99(4): p. 277–285.

35. Jang, S., et al., Critical role of c-jun N-terminal protein kinase in promoting mitochondrial dysfunction and acute liver injury. Redox Biol, 2015. 6: p. 552–564.

36. Tang, Y., et al., Effect of alcohol on miR-212 expression in intestinal epithelial cells and its potential role in alcoholic liver disease. Alcohol Clin Exp Res, 2008. 32(2): p. 355–64.

37. Nannelli, G., et al., ALDH2 Activity Reduces Mitochondrial Oxygen Reserve Capacity in Endothelial Cells and Induces Senescence Properties. Oxidative Medicine and Cellular Longevity, 2018. 2018: p. 9765027.

38. Kang, P., et al., Activation of ALDH2 attenuates high glucose induced rat cardiomyocyte fibrosis and necroptosis. Free Radic Biol Med, 2020. 146: p. 198–210.

39. Chen, C.H., et al., Activation of aldehyde dehydrogenase-2 reduces ischemic damage to the heart. Science, 2008. 321(5895): p. 1493–5.

40. Zhang, T., et al., Alda-1, an ALDH2 activator, protects against hepatic ischemia/reperfusion injury in rats via inhibition of oxidative stress. Free Radical Research, 2018. 52(6): p. 629–638.

41. Chen, L., et al., Vinyl chloride-induced interaction of nonalcoholic and toxicant-associated steatohepatitis: Protection by the ALDH2 activator Alda-1. Redox Biology, 2019. 24: p. 101205.

42. Chaudhry, K.K., et al., ALDH2 Deficiency Promotes Ethanol-Induced Gut Barrier Dysfunction and Fatty Liver in Mice. Alcohol Clin Exp Res, 2015. 39(8): p. 1465–75.

43. Cho, Y.E. and B.J. Song, Pomegranate prevents binge alcohol-induced gut leakiness and hepatic inflammation by suppressing oxidative and nitrative stress. Redox Biol, 2018. 18: p. 266–278.

44. Keshavarzian, A., et al., Evidence that chronic alcohol exposure promotes intestinal oxidative stress, intestinal hyperpermeability and endotoxemia prior to development of alcoholic steatohepatitis in rats. Journal of Hepatology, 2009. 50(3): p. 538–547.

45. Szabo, G. and S. Bala, Alcoholic liver disease and the gut-liver axis. World J Gastroenterol, 2010. 16(11): p. 1321–9.

46. NIAAA, Alcohol Facts and Statistics. National Institute on Alcohol Abuse and Alcoholism, 2021: p. https://www.niaaa.nih.gov/publications/brochures-and-fact-sheets/alcohol-facts-and-statistics.

47. Liangpunsakul, S., P. Haber, and G.W. McCaughan, Alcoholic Liver Disease in Asia, Europe, and North America. Gastroenterology, 2016. 150(8): p. 1786–1797.

48. CDC, Excessive Drinking is Draining the U.S. Economy. Centers for Disease Control and Prevention, 2019: p. https://www.cdc.gov/alcohol/features/excessive-drinking.html.

49. Wu, D. and A.I. Cederbaum, Alcohol, oxidative stress, and free radical damage. Alcohol Res Health, 2003. 27(4): p. 277–84.

50. Lakshman, M.R., et al., CYP2E1, oxidative stress, post-translational modifications and lipid metabolism. Subcell Biochem, 2013. 67: p. 199–233.

51. Rungratanawanich, W., M. Memo, and D. Uberti, Redox Homeostasis and Natural Dietary Compounds: Focusing on Antioxidants of Rice (Oryza sativa L.). Nutrients, 2018. 10(11).

52. Brooks, P.J., et al., The alcohol flushing response: an unrecognized risk factor for esophageal cancer from alcohol consumption. PLoS Med, 2009. 6(3): p. e50.

53. Yokoyama, A., et al., Alcohol-related cancers and aldehyde dehydrogenase-2 in Japanese alcoholics. Carcinogenesis, 1998. 19(8): p. 1383–1387.

54. Yokoyama, T., et al., Alcohol Flushing, Alcohol and Aldehyde Dehydrogenase Genotypes, and Risk for Esophageal Squamous Cell Carcinoma in Japanese Men. Cancer Epidemiology Biomarkers &amp;amp; Prevention, 2003. 12(11): p. 1227.

55. Yu, H.S., et al., Characteristics of aldehyde dehydrogenase 2 (Aldh2) knockout mice. Toxicol Mech Methods, 2009. 19(9): p. 535–40.

56. Liao, J., et al., Aldehyde dehydrogenase-2 deficiency aggravates cardiac dysfunction elicited by endoplasmic reticulum stress induction. Mol Med, 2012. 18(1): p. 785–93.

57. Yukawa, Y., et al., Impairment of aldehyde dehydrogenase 2 increases accumulation of acetaldehyde-derived DNA damage in the esophagus after ethanol ingestion. Am J Cancer Res, 2014. 4(3): p. 279–84.

58. Han, H., et al., Protective Effects of Facilitated Removal of Blood Alcohol and Acetaldehyde Against Liver Injury in Animal Models Fed Alcohol and Anti-HIV Drugs. Alcohol Clin Exp Res, 2019. 43(6): p. 1091–1102.

59. Kim, Y.D., et al., Expression levels of hepatic cytochrome P450 enzymes in Aldh2-deficient mice following ethanol exposure: a pilot study. Arch Toxicol, 2005. 79(4): p. 192–5.

60. Oyama, T., et al., Tissue-distribution of aldehyde dehydrogenase 2 and effects of the ALDH2 gene-disruption on the expression of enzymes involved in alcohol metabolism. Front Biosci, 2005. 10: p. 951–60.

61. Lu, Y. and A.I. Cederbaum, Cytochrome P450s and Alcoholic Liver Disease. Curr Pharm Des, 2018. 24(14): p. 1502–1517.

62. Forsyth, C.B., R.M. Voigt, and A. Keshavarzian, Intestinal CYP2E1: A mediator of alcohol-induced gut leakiness. Redox Biol, 2014. 3: p. 40–6.

63. de Timary, P., et al., A role for the peripheral immune system in the development of alcohol use disorders? Neuropharmacology, 2017. 122: p. 148–160.

64. Rao, R., Endotoxemia and gut barrier dysfunction in alcoholic liver disease. Hepatology, 2009. 50(2): p. 638–644.

65. Chidambaram, S.B., et al., Gut dysbiosis, defective autophagy and altered immune responses in neurodegenerative diseases: Tales of a vicious cycle. Pharmacol Ther, 2021: p. 107988.

66. Rahman, K., et al., Loss of Junctional Adhesion Molecule A Promotes Severe Steatohepatitis in Mice on a Diet High in Saturated Fat, Fructose, and Cholesterol. Gastroenterology, 2016. 151(4): p. 733–746.e12.

67. Abdelmegeed, M.A., et al., Cytochrome P450-2E1 promotes fast food-mediated hepatic fibrosis. Sci Rep, 2017. 7: p. 39764.

68. Frazier, T.H., J.K. DiBaise, and C.J. McClain, Gut Microbiota, Intestinal Permeability, Obesity-Induced Inflammation, and Liver Injury. Journal of Parenteral and Enteral Nutrition, 2011. 35(5S): p. 14S–20S.

69. Sandler, N.G. and D.C. Douek, Microbial translocation in HIV infection: causes, consequences and treatment opportunities. Nature Reviews Microbiology, 2012. 10(9): p. 655–666.

70. Samuelson, D.R., et al., Intestinal Microbial Products From Alcohol-Fed Mice Contribute to Intestinal Permeability and Peripheral Immune Activation. Alcohol Clin Exp Res, 2019. 43(10): p. 2122–2133.

71. Long, E.K., D.M. Olson, and D.A. Bernlohr, High-fat diet induces changes in adipose tissue trans-4-oxo-2-nonenal and trans-4-hydroxy-2-nonenal levels in a depot-specific manner. Free Radic Biol Med, 2013. 63: p. 390–8.

72. Wu, P., et al., Serum TNF-α, GTH and MDA of high-fat diet-induced obesity and obesity resistant rats. Saudi Pharm J, 2016. 24(3): p. 333–6.

73. Anderson, J.M. and C.M. Van Itallie, Physiology and function of the tight junction. Cold Spring Harb Perspect Biol, 2009. 1(2): p. a002584.

74. Rao, R.K., Acetaldehyde-induced barrier disruption and paracellular permeability in Caco-2 cell monolayer. Methods Mol Biol, 2008. 447: p. 171–83.

75. Wang, Z., et al., Increased expression of microRNA-378a-5p in acute ethanol exposure of rat cardiomyocytes. Cell Stress and Chaperones, 2017. 22(2): p. 245–252.

76. Zhu, Q., et al., Pretreatment with the ALDH2 agonist Alda-1 reduces intestinal injury induced by ischaemia and reperfusion in mice. Clin Sci (Lond), 2017. 131(11): p. 1123–1136.

77. Panisello-Roselló, A., et al., Role of aldehyde dehydrogenase 2 in ischemia reperfusion injury: An update. World J Gastroenterol, 2018. 24(27): p. 2984–2994.

